# Biophysical assay for tethered signaling reactions reveals tether-controlled activity for the phosphatase SHP-1

**DOI:** 10.1101/063776

**Authors:** Jesse Goyette, Citlali Solis Salas, Nicola Coker-Gordon, Marcus Bridge, Samuel A. Isaacson, Jun Allard, Omer Dushek

## Abstract

Tethered enzymatic reactions are ubiquitous in signalling networks but are poorly understood. Here, a novel mathematical analysis is established for tethered signalling reactions in surface plasmon resonance (SPR). Applying the method to the phosphatase SHP-1 interacting with a phosphorylated tether corresponding to an immune receptor cytoplasmic tail provides 5 biophysical/biochemical constants from a single SPR experiment: two binding rates, two catalytic rates, and a reach parameter. Tether binding increased the activity of SHP-1 by 900-fold through a binding-induced allosteric activation (20-fold) and a more significant increase in local substrate concentration (45-fold). The reach parameter indicates that this local substrate concentration is exquisitely sensitive to receptor clustering. We further show that truncation of the tether leads not only to a lower reach but also to lower binding and catalysis. The work establishes a new framework for studying tethered signalling processes and highlights the tether as a control parameter in clustered signalling.

## Introduction

A common theme in signal transduction pathways is the tethering of signalling enzymes near their substrates before catalysis (1, 2). Familiar examples include reactions on surface receptors, where cytoplasmic enzymes first bind to receptor tails (tethers) before catalysing reactions on substrates within reach. Understanding these complicated reactions is limited because they depend not only on the catalytic rate but also on the binding kinetics that localise the enzyme and on the tether reach. Moreover, many cell surface receptors are known to cluster but how clustering influences reaction rates is poorly understood (3).

A large group of immune surface receptors rely on the tethering of cytoplasmic kinases and phosphatases to both initiate and integrate signalling (4). Their unstructured cytoplasmic tails contain multiple tyrosines that serve as both docking sites and substrates for these enzymes. In the case of inhibitory immune receptors (e.g. PD-1, LAIR-1), tyrosines in conserved immunotyrosine-based inhibitory or switch motifs (ITIMs or ITSMs) generate docking sites for the SH2 domains of the cytosolic phosphatases SHP-1 and/or SHP-2. When tethered, these phosphatases are thought to undergo allosteric catalytic activation (5–8) to dephosphorylate various membrane-proximal tyrosines (9–11).

Microscopy studies have highlighted the clustering of immune receptors on the plasma membrane (11–14) but the consequences of clustering remain poorly defined. For example, it is presently unknown how membrane-localisation and the degree of clustering influences the local substrate concentration experienced by SHP-1. Mathematical models can predict large local substrate concentrations for certain tethers (15–17), which may even override the catalytic specificity of the enzyme (18). This may explain the observation that SHP-1 and SHP-2 can regulate the phosphorylation state of the clustered inhibitory receptors they interact with (11).

Solution-based *in vitro* assays for enzymatic activity have been instrumental to our understanding of signalling, and in particular to SHP-1 (5,6). These experiments measure the reaction product over time after mixing the enzyme and substrate in solution and model fitting produces an estimate for the overall catalytic rate 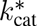(= *k_cat_/K*_m_, where *K*_m_ is the Michaelis constant) (19). Applying this assay to SHP-1 (Fig. 1a-b) makes it clear that this single number coarse-grains the reaction mechanism when proteins have multiple domains that interact with substrates and moreover, the tether does not influence these reaction rates.

Surface plasmon resonance (SPR), as implemented in commercial instruments such as BIAcore™ (GE Healthcare), is a widely used biophysical assay for molecular interactions (20). In a typical experiment, one binding partner is immobilised to a surface while the other is injected over it. The instrument reports a high accuracy measure of the mass of material bound at the surface, expressed as Resonance Units (RU), as a function of time. The resulting data is fit to mathematical models to determine the association rate (*k*_on_) and dissociation rate (*k*_off_). High sensitivity, accuracy, and the availability of many surface chemistries have meant that the method has gained considerable popularity not only for biomedical research but for medical diagnostics, food safety and security, and environmental monitoring (20). Despite these advances the method remains largely a tool for the study of molecular binding.

In this work we retool SPR for the study of tethered enzymatic reactions applied to SHP-1. Injection of SHP-1 over a surface immobilised with phosphorylated peptides produced a non-canonical SPR trace as a result of tethered dephosphorylation reactions of clustered peptides. Using a novel mathematical analysis, we show that 5 biophysical and biochemical constants can be independently extracted from a single SPR trace. We found that binding of either SH2 domain to the tether allosterically activated SHP-1 and that the tether length modulated not only the reach but also binding and catalysis. Using these parameters we find that tethering increases reaction rates by 900-fold with a tether-induced local increase in substrate concentration the dominant contribution but only when receptors are clustered within 5 nm. Collectively, the work highlights tethering as a control parameter for signalling reactions and provides a novel SPR-based platform for the study of tethered signalling, with implications to drug discovery.

**Figure 1:**
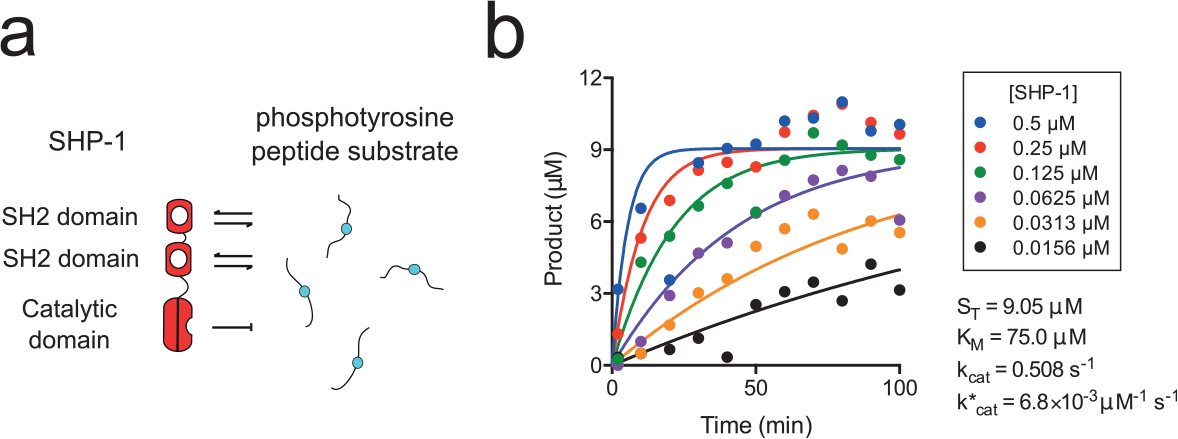
Solution assay for SHP-1.(**a**) Schematic of the domain structure of SHP-1 and a subset of reactions that may occur in solution with peptide substrates. (**b**) Standard solution-based enzymatic assay showing the production of inorganic phosphate (product) over time for the indicated concentration of SHP-1 mixed with PEG12-ITIM substrate (data is representative of two independent experiments). Progress curves are fit with a mathematical model to provide an estimate for 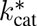 (see Materials & Methods).

## Results

### Non-canonical SPR traces as a result of tethered enzymatic reactions

To create a substrate surface for SHP-1 in SPR, we coated a surface with peptides containing an ITIM sequence, from the N-terminus of the inhibitory receptor LAIR-1, with a 28-repeat polyethylene glycol (PEG) spacer (PEG28-ITIM, Table 1). When SHP-1 was injected over this surface we observed a complicated curve with an initial binding phase that was quickly followed by a reduction in binding despite continuous injection of SHP-1 (Fig. 2a). To convert the arbitrary response units reported by the SPR instrument to a more meaningful unit we normalised this curve to maximum binding assuming 1-to-1 interaction with peptide (see Materials & Methods). We confirmed that SHP-1 was dephosphorylating the substrate by injecting an anti-phosphotyrosine antibody following the injection of SHP-1, which revealed near complete dephosphorylation (Fig. 2b).

The decrease in binding, despite continuous injection of SHP-1, can be understood by considering the catalytic activity of the enzyme that over time destroys binding sites for its SH2 domains. When SHP-1 is first injected over the surface it begins binding via the SH2 domains (initial rise within the first 3 s in Fig. 2a). This binding (or tethering) increases the dephosphorylation rate by confining SHP-1 and its phosphorylated substrates to a restricted volume resulting in the rapid destruction of highly clustered phosphorylated peptides (steep fall between 3 and 100 s in Fig. 2a). However, the rate of dephosphorylation by tethered SHP-1 decreases over time because the tethered enzyme is unable to reach remaining phosphorylated peptides, whose average distance increases over time. This inefficient tethered dephosphorylation combined with inefficient solution dephosphorylation leads to a slow loss in overall binding at later timepoints (slowly decreasing asymptote after 100 s in Fig. 2a). These interactions are summarised in Fig. 2c.

**Figure 2:**
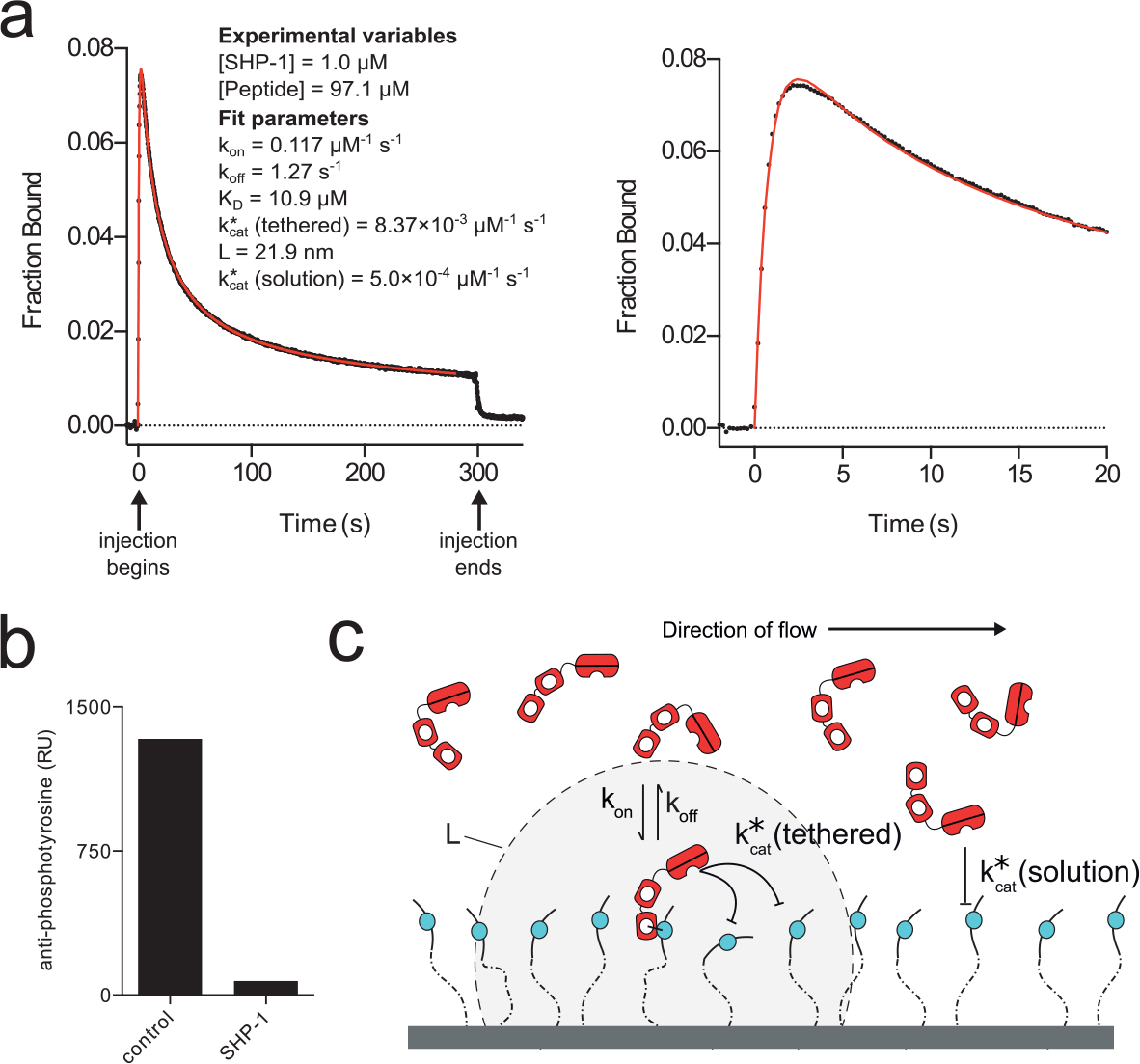
A tethered enzymatic surface plasmon resonance assay for SHP-1. (**a**) Representative SPR trace (black dots) for SHP-1 injected over a surface immobilised with 48.5 RU of an ITIM peptide derived from LAIR-1 on a 28 repeat polyethylene glycol linker (PEG28-ITIM). A fit of the MPDPDE model (red line) provides estimates for the indicated parameters. Early time data and fit is shown on the right. Unprocessed data is shown in Supplementary Fig. 1. (**b**) Anti-phosphotyrosine antibody injected at the end of the experiment shows reduced binding in the experimental flow cell (SHP-1) compared to a buffer-injected flow cell with equivalent peptide levels (control). (**c**) Schematic of reactions taking place when SHP-1 is injected over a surface of immobilised phosphorylated peptides. Note that peptide anchoring is displayed in 1D for clarity but because the surface consists of a dextran matrix onto which peptides randomly couple they are anchored in 3D.

### A mathematical model quantifies the tethered enzymatic SPR assay

The tethered enzymatic SPR assay is heavily influenced by stochastic fluctuations. This may seem counter-intuitive because the instrument reports macroscopic binding averaged over picomoles of protein across a millimeter scale surface. However, tethered catalytic reactions are limited to the number of peptide substrates within reach, which we estimate to be ~8 initially (assuming [Peptide] = 100 *μ*M with a reach of 25 nm) and over time to reach 0. Therefore, the SPR trace represents the average of many realisations of a low copy number stochastic process.

We therefore developed a spatial stochastic simulation to reproduce the tethered enzymatic SPR assay. The model includes: the kinetics of SHP-1 binding to phosphorylated peptides by its SH2 domains (governed by on-rate, *k*_on_, and off-rate, *k*_off_, constants), dephosphorylation of peptides when SHP-1 is bound to the surface, 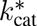 (tethered), or when SHP-1 is free in solution, 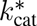 (solution) (Fig. 2c). The local concentration of peptide experienced by tethered SHP-1 is determined by the reach parameter *L*, which is defined as the quadrature average of the average reach distance of the peptide and the average reach distance of SHP-1 bound to a peptide. This calculation is based on approximating the motion of both the free and bound peptide using the worm-like-chain polymer model (see Materials & Methods). The stochastic simulation was used to plot the three molecular species over time and to provide spatial snapshots of these species at different times (Fig. 3). As expected, we found that clustered peptides were preferentially destroyed leading to a non-random distribution of the surviving phosphorylated peptides.

**Figure 3:**
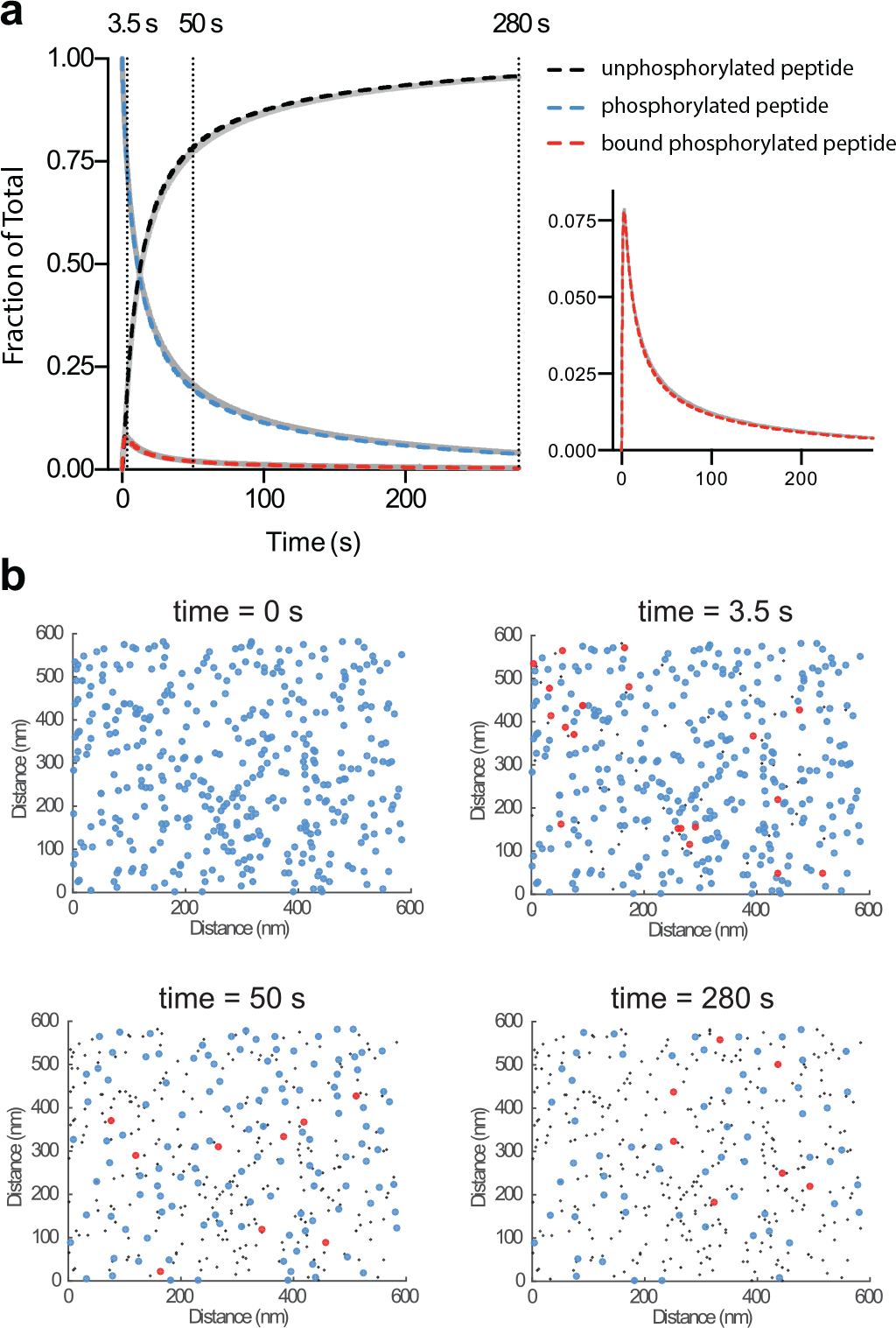
Mathematical models capture the physics and chemistry in tethered enzymatic SPR. (**a**) Levels of unphosphorylated peptide, phosphorylated peptide, and phosphorylated peptide bound to SHP-1 over time determined using the stochastic simulation (solid grey lines) or the MPDPDE model (dashed coloured lines). Levels of SHP-1 bound to phosphorylated peptide is re-plotted for clarity (right panel). Good agreement is observed between the stochastic simulation and the MPDPDE model calculation. (**b**) Snapshots of the spatial distribution of the 3 molecular species at indicated timepoints from the stochastic simulation (colours as in panel a). Initially, the phosphorylated peptides are randomly distributed on the surface (0 s), but as time progresses clustered peptides are effectively dephosphorylated by tethered catalysis (50 s) ultimately resulting in phosphorylated peptides too far apart for efficient tethered catalysis (280 s). These 2D spatial distributions are generated from the stochastic simulation by projecting 20 nm in the third dimension. See Materials & Methods for computational details. Parameters: [SHP-1] = 1 *μ*M, [Peptide] = 100 *μ*M, *k*_on_ = 0.1 *μ*M^−1^s^−1^, *k*_off_ = 1 s^−1^, 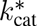 (tethered) = 0.01 *μ*M^−1^s^−1^, L = 20 nm, and 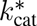(solution) = 0.0005 *μ*M^−1^s^−1^.

Stochastic simulations often provide intuition but are not practical for data fitting because they require long computation times. We therefore developed a computationally efficient model. Models based on standard partial-differential-equations (PDEs) fail to fit the experimental data because they do not account for stochastic fluctuations (not shown). We therefore used the multi-particle density (MPD) formalism, previously used to study defects in solid state physics (21–23), to develop a hybrid integral MPDPDE model that includes the reactions specified for the stochastic simulation (see Materials & Methods). We found good agreement between the stochastic simulation and the computationally efficient MPDPDE model (Fig. 3a).

We used the MPDPDE model to examine the dependency of the predicted SPR trace on the experimental variables (SHP-1 and peptide concentrations) and on the 5 model parameters (*k*_on_, *k*_off_, 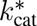 (tethered), *L*, and 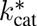 (solution)). We found that the binding trace shifted in non-intuitive ways (Supplementary Fig. 2). For example, changing the concentration of peptide, which in standard SPR simply changes the scale of the binding trace, resulted in a change to the shape of the binding trace because a different proportion of peptides were dephosphorylated by tethered versus solution enzyme.

We next analysed the SPR data using the MPDPDE model. We found an excellent fit of the model to the data (Fig. 2a, red line) and recovered the 5 model parameters. We performed Markov-Chain Monte-Carlo analysis to determine whether a different set of parameters can produce the same binding trace but found that the 5 recovered parameters are unique (Supplementary Fig. 3). In summary, the computationally efficient MPDPDE model captures the stochastic fluctuations in tethered reactions and is able to recover 5 parameters from a single SPR trace.

### Fitted biophysical and biochemical constants are independent of experimental variables

A key test of a mathematical model is the ability to recover the same parameter values when different experimental variables are used. This is particularly important for SPR where mass transport and rebinding can produce parameters that are dependent on the concentration of surface immobilised receptor (24, 25). We therefore performed experiments at different SHP-1 concentrations and immobilised peptide concentrations (Fig. 4a,b). The fitted parameters did not correlate with either experimental variable and moreover, showed excellent reproducibility (Fig. 4c). As predicted by the model, changing the concentration of immobilised peptide led to a change in the shape of the SPR binding trace (Fig. 4b), highlighting the difficulty of interpreting the data without performing model fitting.

### Fitted biophysical and biochemical constants are consistent with the biology of SHP-1

The recovered parameters (Fig. 4c) were within the expected range for tethered signalling by SHP-1. The affinity of SHP-1 interacting with the LAIR-1 ITIM (*K*_D_ = 9.38 *μ*M) is in good agreement with those for isolated SH2 domains of SHP-1 interacting with other ITIMs (26). We observed a 20-fold increase in the tethered over solution catalytic rate (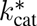 (tethered) =1.18×10^−2^ *μ*M^−1^s^−1^ vs. 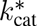 (solution) =6.03×10^−4^ *μ*M^−1^s^−1^), which is consistent with an allosteric activation of SHP-1 when bound by SH2 domains (6). We note that the standard solution-based assay recovered an overall catalytic rate that was between these two rates (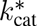 =6.8×10^−3^ *μ*M^−1^s^−1^, Fig. 1b) consistent with a combination of inactive and allosterically active SHP-1 mediating dephosphorylation in solution.

The tethered enzymatic SPR assay also produced an estimate for the reach parameter (*L*), which was found to be 23 nm. This number corresponds to a phosphorylated substrate experiencing a maximum local SHP-1 concentration of 45 *μ*M when tethered, compared to, for example, a concentration of 1 *μ*M in solution (Fig. 2a) or in the cytoplasm of immune cells (see Discussion). To further appreciate the effect of surface tethering, we used the fitted parameters to calculate the time required to dephosphorylate 50% of the peptides with tethering (9.2 s) and without tethering (19 min), revealing that tethering reduced the reaction time by 125-fold (Supplementary Fig. 5).

**Figure 4:**
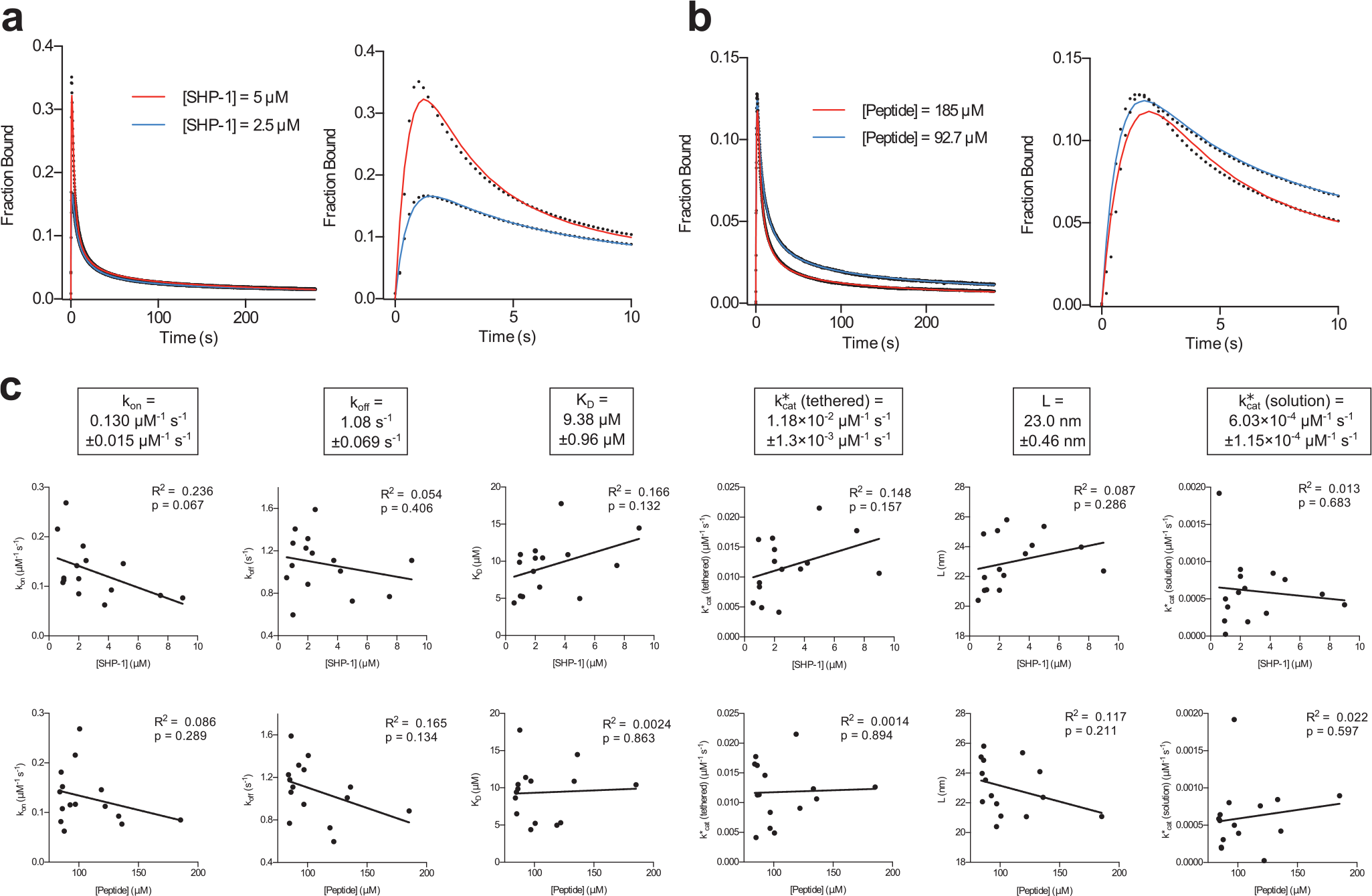
Fitted parameters are independent of experimental variables. Representative SPR traces (black dots) and MPDPDE model fits (solid colour) for (**a**) two SHP-1 concentrations and (**b**) two initial peptide concentrations with right panels showing early time data and fit. (**c**) Plots of fitted parameters versus SHP-1 concentration (upper row) and peptide concentration (lower row) with linear regression fits (R^2^ and p values are indicated) reveal a lack of correlation, indicating that the fitted parameters do not depend on the experimental variables. Averages of fit parameters with SEMs from all experiments are shown in boxes (n=15). All experiments are performed using wild-type SHP-1 and phosphorylated PEG28-ITIM peptides. Exclusion criteria for experiments exhibiting long timescale artefacts, such as non-specific binding and/or differential flow cell drift, are discussed in the Materials & Methods (Quality Control) and Supplementary Fig. 4.

### The tether length controls the binding, catalysis, and reach parameters

To further understand the effects of the reach parameter (*L*) we performed experiments with different tether lengths. To do this, we reduced the length of the spacer from 28 to 0 PEG repeats without modifications to the peptide (Table 1). The model produced excellent fits to all data (Fig. 5a) and, as expected, the reach parameter decreased with decreasing tether lengths (Fig. 5b). Interestingly, although there is a large difference in the contour length between the longest (PEG28, 12.1 nm) and shortest (PEG0, 2.3 nm) tethers, there was not a dramatic decrease in *L* (23 nm for PEG28 and 17 nm for PEG0). This implied that SHP-1 itself contributes significantly to the reach length when it is tethered. This can be understood by noting that, although the contour length of the tethers may be long, the average reach distance is relatively short due to the small persistence length of flexible PEG (27) and polypeptides (28). Thus the rigid domains of SHP-1 may contribute much more to the reach length than one might intuitively expect.

We note that a decrease in *k*_on_ and 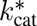 (tethered) were also observed (Fig. 5b), which likely reflects steric hindrance at short tether lengths (PEG3 and PEG0). This form of configurational hindrance is a result of a smaller fraction of time that short linkers spend sufficiently far from their anchor point to accommodate SHP-1 binding. This mechanism is not expected to change *k*_off_, which is consistent with the similar *k*_off_ values we find across tethers. This effect is significant and indicates that SPR experiments to measure binding to phosphorylated peptides should utilise long PEG linkers to avoid configurational steric hinderance.

**Figure 5:**
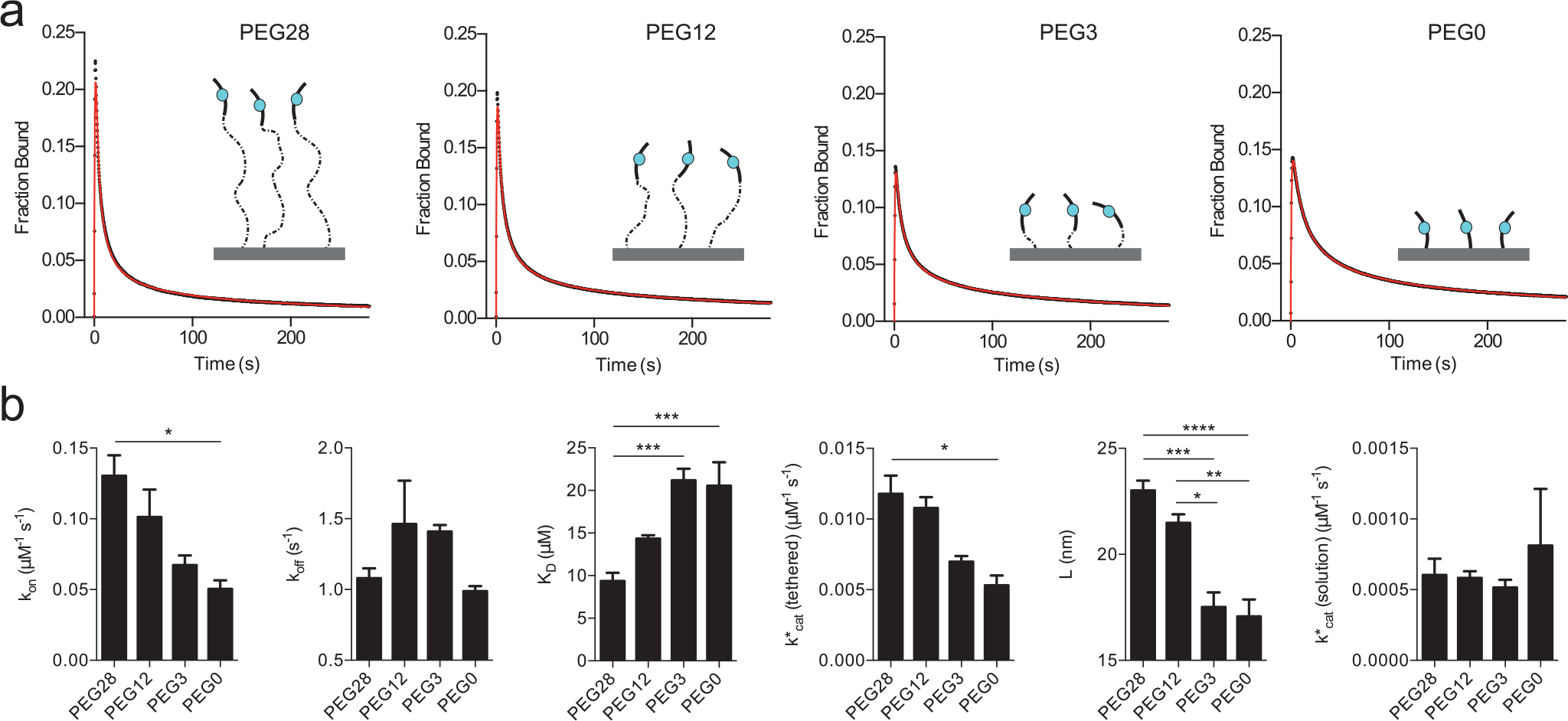
Reduction in tether lengths reduces the reach parameter and introduce configurational steric hindrance. (**a**) SPR traces (black circles) and MPDPDE model fits (red lines) of SHP-1 injected over peptides with 28 (PEG28), 12 (PEG12), 3 (PEG3) and 0 (PEG0) PEG linker repeats. (**b**) Average fit parameters for PEG28-ITIM (n=15), PEG12-ITIM (n=3), PEG3-ITIM (n=3) and PEG0-ITIM (n=2) show reduced values of *L*, *k*_on_, and 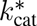 (tethered) for shorter linkers. Two-way ANOVA with a Bonferroni multiple comparison correction is used to determine p values (**** p< 0.0001, *** p< 0.001, ** p< 0.01, * p< 0.05).

### A different reach but a similar allosteric activation is induced by each SH2 domain of SHP-1

Given that SHP-1 itself may significantly contribute to the reach length, we hypothesised that binding by the N-terminal SH2 domain would allow SHP-1 to reach further compared to the C-terminal SH2 domain. We generated SHP-1 variants with inactivating point mutations to either SH2 domain. Mutation of the N-terminal SH2 domain showed drastically reduced binding, whereas mutation of the C-terminal SH2 domain had a weak effect on binding (Fig 6a-b), clearly demonstrating that the N-terminal SH2 domain dominates the interaction of wild type SHP-1 to the membrane proximal ITIM of LAIR-1.

As expected, a reduction in the reach parameter was observed for the N-terminal mutant (*L* = 16.5 nm) compared to the wild-type (*L* = 23.0 nm) and C-terminal mutant (*L* = 23.9 nm) because binding via the C-terminal SH2 reduced the overall reach (Fig. 6b). This difference is more than twice as large as the spatial extent of the C-terminal SH2 domain (~3 nm, estimated from structure), which reflects the large effective persistence length of structured domains.

Interestingly, and in contrast to previous studies, we found that the N-terminal mutant still exhibited allosteric activation because 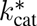 (tethered) remained 10-fold larger than 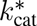 (solution). Therefore, binding of either the N or C-terminal SH2 domains is sufficient to allosterically activate SHP-1. We also find that the *k*_on_ differed by ~10-fold between the N-and C-terminal SH2 domains but the *k*_off_ was nearly identical (Fig. 6b).

**Figure 6:**
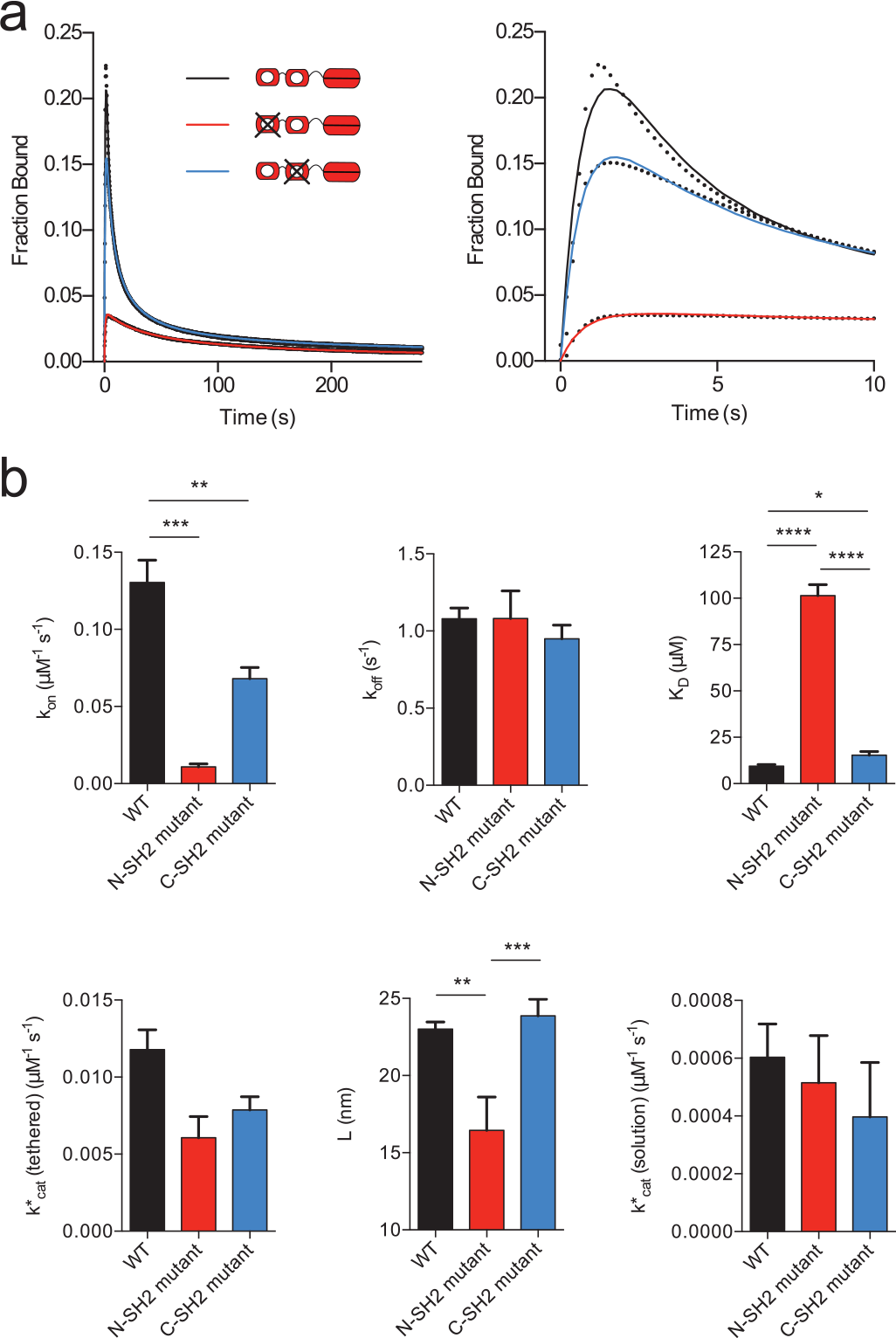
Binding by either SH2 domain allosterically activates SHP-1 but the reach parameter is larger for N-SH2 binding. (**a**) SPR traces (black dots) and MPDPDE model fits (solid lines) of N-terminal and C-terminal SH2 domain binding-null mutants and wild-type SHP-1 injected over PEG28-ITIM. (**b**) Average fit parameters for wild-type (WT) SHP-1 (n=15), N-SH2 mutant SHP-1 (n=3), and C-SH2 mutant SHP-1 (n=9) show weak binding and reduced reach when SHP-1 binds via the C-SH2 domain compared to the N-SH2 domain but allosteric activation is observed in both cases. Two-way ANOVA with a Bonferroni multiple comparison correction is used to determine p values (**** p< 0.0001, *** p< 0.001, ** p< 0.01, * p< 0.05)

## Discussion

Our understanding of tethered signalling reactions is limited by a lack of experimental methods. We have described a novel SPR-based assay for tethered enzymatic reactions that, from a single experiment, can recover 5 biophysical and biochemical constants that quantify tethered signalling for SHP-1 with clustered substrates. We demonstrate that these constants can be determined with high accuracy, as a result of the high sensitivity of SPR, and further show that they are independent of the SHP-1 and substrate concentrations.

We observed that reducing tether lengths below ≈12 nm (PEG28) introduces a steric penalty to binding, implying a lower bound on the cytoplasmic tails of inhibitory receptors that recruit SHP-1 (Fig. 7a). Indeed, a bioinformatic analysis of inhibitory receptors reveals that the majority of receptors contain ITIMs that are ≥12 nm from the plasma membrane (Fig. 7b). Interestingly, most activatory receptors contain tyrosines that are located ≤12 nm from the plasma membrane. This finding raises the possibility that tethers may have a role in binding specificity by, for example, sterically preventing binding of signalling enzymes.

Activatory and inhibitory immune receptors are both known to cluster in the plasma membrane (11–14) but the extent and consequences of clustering remain poorly understood. In the absence of clustering, a phosphorylation site on the cytoplasmic tails of these receptors will experience the low activity of cytoplasmic SHP-1 at a concentration of ~ 1 *μ*M (based on 280,000 copies of SHP-1 in cytotoxic T cells (29) of radius 5 *μ*m). Tethering of SHP-1 to non-clustered immune receptors at distances > 50*nm* results in low SHP-1 concentrations (e.g. 0.04 *μ*M when receptors are 50 nm apart) but when clustered within 5 nm, we can now estimate that this phosphorylated site will experience a SHP-1 concentration of 45 *μ*M (Fig. 7c). This concentration is exquisitely sensitive to the degree of clustering so that a 10-fold decrease in receptor clustering (5 to 50 nm) results in a 1125-fold decrease in concentration (45 to 0.04 *μ*M). These concentrations are calculated using the formula for *σ* with *L* = 23 nm for *r* = 50 nm or *r* = 5 nm (see Materials & Methods). The 45-fold increase in SHP-1 concentration is based on a reach of *L* = 23 nm, which represents SHP-1 bound to an ITIM on PEG28 dephosphorylating another ITIM on PEG28. We expect the value of *L* to decrease, and hence the local concentration to increase, when SHP-1 dephosphorylates other substrates such as ITAMs on shorter activatory receptor tails.

The current model for SHP-1 activation is based on an allosteric conformational change into an ‘open’ high catalytic activity state induced by N-terminal SH2 domain binding (6, 8, 30). In agreement with this model, we have found a 20-fold increase in the catalytic rate when the SH2 domains are engaged (tethered vs solution catalytic rates, Fig. 4c). The sensitivity of the present assay has revealed that the C-terminal SH2 domain is also able to induce the open state, suggesting that the ‘closed’ low activity state may involve occlusion of the catalytic domain by either SH2 domain. The observation that unbound SHP-1 exhibits catalytic activity, albeit less efficient, suggests that it is in equilibrium between closed and open states in solution. Assuming the open state transition is complete upon SH2 binding, and that the activity of the open state is similar when tethered or when achieved spontaneously when unbound, our results suggest that SHP-1 spends only 5% of the time in the open active conformation in solution (i.e. 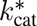 (solution) = 0.05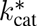 (tethered)).

The combined effects induced by SHP-1 tethering on allosteric activation and local concentration can be summarised by calculating the overall dephosphorylation rate (Fig. 7c). We find a similar rate when SHP-1 is in solution or tethered but not clustered (1 *μ*M × 0.000603 *μ*M^−1^s^−1^ vs. 0.04 *μ*M × 0.0118 *μ*M^−1^s^−1^) but observe a 900-fold increase in the dephosphorylation rate when tethered and clustered (45 *μ*M × 0.0118 *μ*M^−1^s^−1^).

The biophysical assay for tethered enzymatic reactions introduced here can be used for the study a large number of tethered signalling reactions on immune receptors (4). Although we have focused on the interactions with the tyrosine phosphatase SHP-1, the assay can be performed with the large number of tyrosine kinases, such as those of the Src-and Syk-families, that can both phosphorylate and bind their substrates. More generally, the method can be used in any situation where an enzyme can both bind and modify a substrate. Many such enzymes, SHP-1/SHP-2 included, are attractive therapeutic targets and by providing rich mechanistic information, the assay may be particularly useful to identify drugs that target allosteric mechanisms (31) or tether components. Unlike the catalytic domains of the enzymes they recruit, tethers such as immune receptor cytoplasmic tails are often conserved in length but not sequence, potentially allowing for more targeted therapeutics. The tethered enzymatic assay is a useful extension to the already widely used SPR platform for drug discovery and mechanistic studies (20) but we expect that it can be implemented in other instruments where binding can be observed over time (e.g. Bio-Layer Interferometry).

**Figure 7:**
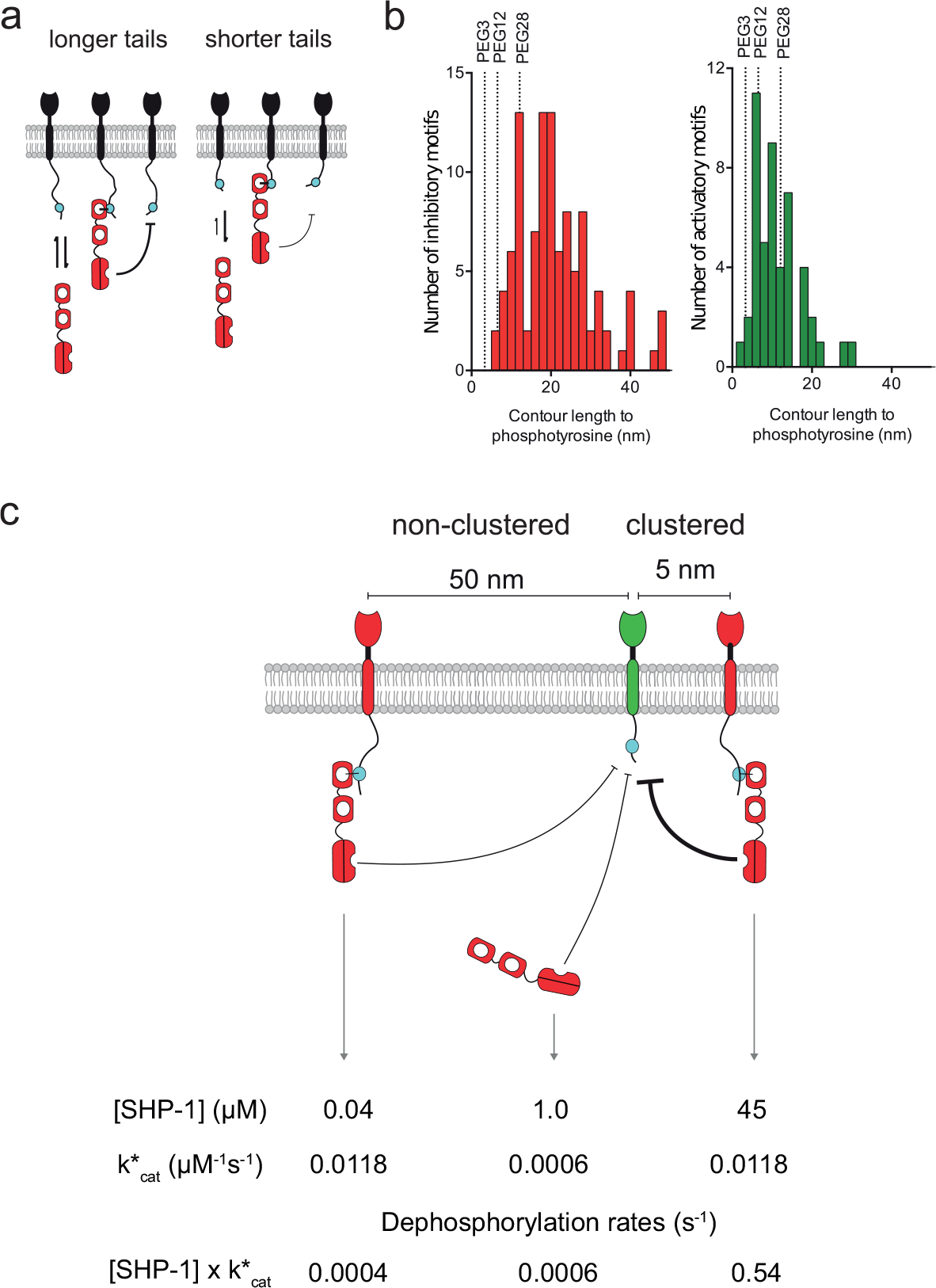
Diagramatic summary. (**a**) Shorter cytoplasmic tails (tethers) result in decreased *k*_on_ for binding and 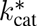 (tethered) for catalysis as a result of steric hindrance (indicated by thickness of black arrow). (**b**) Histograms of the contour length (number of amino acids × 0.3 nm per amino acid) to phosphotyrosine residues on inhibitory receptors (ITIM/ITSM, mean length of 21.2 nm, red histogram) and activatory receptors (ITAM/ITTM/YxxM, mean length of 11.3 nm, green histogram) for human receptors. The majority of inhibitory receptor tails are longer than PEG28 whereas the majority of activatory receptor tails are shorter than PEG28. (**c**) Overall dephosphorylation rates for SHP-1 tethered to non-clustered receptors (left), free in the cytoplasm (centre), or tethered to clustered receptors (right). Dephosphorylation rates are calculated by multiplying the SHP-1 concentration experienced by the substrate with the catalytic rate of each state. We find that the dephosphorylation rate is 900-fold larger (0.54/0.0006) when SHP-1 is tethered in clustered receptors as a result of a combination of allosteric activation (20-fold) and enhanced local concentration (45-fold). Calculation of concentrations and dephosphorylation rates are described in the Discussion.

Tethered signalling reactions are complicated to study because they depend on multiple factors, such as binding kinetics, catalytic rates, allosteric activation, clustering, and tether length/flexibility. The SPR assay for tethered enzymatic reactions is able to recover 5 independent biophysical/biochemical parameters that quantifies these reactions in detail. When applied to SHP-1, the work has revealed that tethering increases enzymatic rates by 900-fold and that this increase is highly sensitive to the degree of receptor clustering. The work provides a new way to quantitatively study tethered signalling processes and has underlined the tether as a control parameter for signalling.

## Materials & Methods

### Plasmids & peptides

A construct expressing murine SHP-1 with an N-terminal 6× His tag was provided by Marion H. Brown. Mutation of the SH2 domains was performed using a quick-change strategy. The mutations introduced were R30K and R33E for the N-terminal SH2 domain mutant and R136K for the C-terminal SH2 domain mutant previously shown to result in loss of binding (6). All peptides were ordered from PeptideSynthetics (Hampshire, UK) and were certified >95% pure. Sequences of peptides used are shown in Table 1. Peptides sequences were derived from the membrane-proximal ITIM sequence of mouse LAIR-1 receptor.

**Table 1:** Peptides used in study (phosphotyrosines are denoted as Y*)

| Name | Sequence | Contour Length |
| --- | --- | --- |
| PEG28-ITIM | biotin-(PEG)<sub>28</sub>-DLQEVTY*IQLDHH | 12.1 nm |
| PEG12-ITIM | biotin-(PEG)<sub>12</sub>-DLQEVTY*IQLDHH | 6.5 nm |
| PEG3-ITIM | biotin-(PEG)<sub>3</sub>-DLQEVTY*IQLDHH | 3.3 nm |
| PEG0-ITIM | biotin-GDLQEVTY*IQLDHH | 2.7 nm |

### Protein production

SHP-1 DNA constructs were transformed into BL21-CodonPlus(DE3)-RIPL strain (Agilent Technologies, Palo Alto, CA, USA) *Escherichia coli* and plated on LB agar with 100 *μ*g/ml ampicillin then grown overnight at 37°C. The next day colonies were innoculated into a 10 ml LB selection media (LB media with 100 *μ*g/ml ampicillin, 50 *μ*g/ml chloramphenicol), grown overnight at 37°C then transferred to 1 L of LB selection media without chloramphenicol until the optical density at 600 nm was 0.6-0.8. The cells were then treated with isopropyl-1-thio-D-galactopyranoside (final concentration 0.1 mM) and harvested by centrifugation after 20 h culture at 25°C.

Bacterial pellets were resuspended in Tris buffered saline (TBS; 20 mM Tris(hydroxymethyl)aminomethane, 150 mM NaCl) with 0.5% Triton X-100 and protease inhibitors (protease inhibitor cocktail, Sigma) then lysed with three 30 second bursts of sonication interspersed with 60 second rest periods on ice. Lysates were clarified by centriguation at 15 000 rcf followed by filtration through a 0.45 *μ*m filter. Clarified lysates were applied to Ni^2+^-NTA resin, which was washed with 10 column volumes of TBS, followed by 10 column volumes of TBS with 30 mM imidizole, before SHP-1 protein was eluted with 50 mM imidizole in TBS with 10%, pH 7.5. Glycerol was added to a final concentration of 10% (v/v) and protein was stored in aliquots at −40°C until the day of experiment.

On the day of the experiment aliquots SHP-1 and mutants were thawed and further separated by size exclusion chromatography Akta FPLC (GE Healthcare Life Sciences) on a Superdex S75 10/300 G column (GE Healthcare Life Sciences) equilibrated with 20 mM HEPES, 150 mM NaCl, 0.05% Tween 20 and 1 mM dithiothreitol. Concentrations of fractions containing SHP-1 were measured using the optical density at 280 nm, using a Nanodrop™ ND1000 spectrophotometer (Thermo Scientific).

### Surface Plasmon Resonance

Experiments were performed on a BIAcore™ 3000 instrument (GE Healthcare Life Sciences). All experiments were performed at 10°C and with a buffer flow-rate of 10 *μl*/min. The buffer used was HEPES buffered saline (HBS-EP, GE Healthcare Life Sciences), which contained: 10 mM HEPES (pH 7.4), 150 mM NaCl, 3 mM EDTA, and 0.005% Surfactant P20.

A CM5 sensorchip was coupled with streptavidin to near saturation (typically between 4000-7000 RU) using the amine coupling kit (GE Healthcare Life Sciences) as described previously(32). After streptavidin was coupled, biotinylated peptides were injected to give the indicated concentrations in experimental flow cells and excess biotin-binding sites were blocked with biotin in HBS-EP. The molar ratio of peptide to streptavidin was kept below 1:1 to ensure peptide immobilisation was random and not clustered on the tetravalent streptavidin molecules. Reference flow cells were treated with buffer then blocked with biotin; pilot experiments using unphosphorylated control peptides in reference flow cells were indistinguishable from buffer-treated reference flow cells when injected with SHP-1. The chip surface was then conditioned with 5× 5 min injections of HBS-EP. SHP-1 protein in HBS-EP with 1 mM dithiothreitol was then injected over reference and experimental flow cells in series for 5 min at the indicated concentrations.

All SPR data was converted from the reference-subtracted data (in RU) to fraction bound by dividing the RU by the theoretical maximum RU expected if the experimental flow cell was saturated with bound SHP-1.

### Determination of peptide concentration

To determine the concentration of peptide in the assays we first needed a conversion factor between RU and mass of peptide at the chip surface. We determined this conversion factor by injecting four concentrations of SHP-1 over a control flow cell on a CM5 chip and plotting the mass of SHP-1 injected against the raw RU change. We repeated this on seven flow cells across four sensor chips to get an average slope of 149±15 RU per g/L (±SEM). This constant, together with the molecular weight of the peptide, was used to convert between the RU of peptide immobilised and the molar concentration at the chip surface. For example, 48.5 RU of PEG28-ITIM (molecular weight 3221) was immobilised to obtain [Peptide] = 97.1 *μ*M in Fig 2a.

### Quality control

From the MPDPDE model-simulated SPR traces one would predict that the 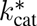 (solution) (Supplementary Fig. 2g,h) and L (Supplementary Fig. 2f) are likely to be very sensitive to small systematic errors in the SPR trace at longer timescales. Non-specific binding and baseline drift are two well-known sources of such systemic errors that can produce artefacts at long timescales (see Supplementary Fig. 4a for examples). To exclude data affected by these artefacts we propose a simple quality control check that greatly improves the accuracy of estimating L, 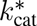 (tethered), and 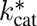 (solution). As a measure of the signal to noise ratio at long time points we took the RU 20 seconds after the injection had completed (noise) and divided it by the RU 20 seconds before the injection finished (signal). We found that when the signal-to-noise ratio was greater than 20% large aberrations in L, 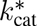 (tethered) or 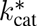 (solution) were apparent, depending on whether the drift was above or below baseline (Supplementary Fig. 4b). Data that had evidence of significant artefact were excluded from the study based on this criteria (red data points in Supplementary Fig. 4b).

### Calculation of local substrate concentration using a polymer model

A key component of the models (described below) is the calculation of the local substrate concentration that a tethered enzyme experiences. We assume that the motion of an unbound phosphorylated peptide (state *A*) and the motion of SHP-1 bound to a phosphorylated peptide (state *B*) can both be approximated by the worm-like-chain model, which is a widely used polymer model. This model provides the probability of finding the tip of the polymer at position **r**,

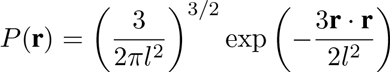

Where 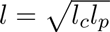 with *l_c_* the contour length and *l_p_* the persistence length. When applied to the free phosphorylated substrate this probability is taken to be the position of the phosphorylated tyrosine residue with *l* = *L_A_.* When applied to bound SHP-1 this probability is taken to be the position of the catalytic pocket of the phosphatase domain with *l* = *L*_*B*_. Using these probabilities, we can calculate the concentration of substrate, *σ*(*r*), that a tethered enzyme will experience when they are anchored a distance of *r* apart (Supplementary Fig. 6),

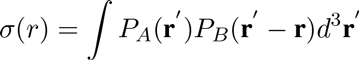

Where the integration is over all space. Without loss of generality we let **r** = *r***ẑ**, which leads to,

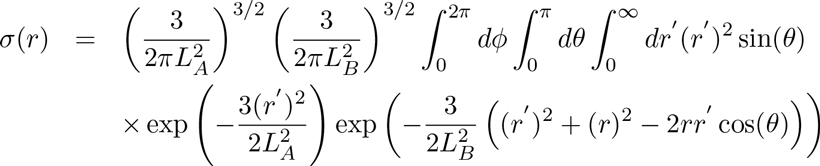

After some algebra, including the variable substitution *q* = (*r*′)^2^ + *r*^2^ – 2*rr*′ cos(*θ*), we find,

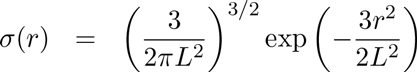

Where 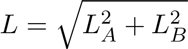 is the reach parameter.

### Mathematical model reactions

We developed a stochastic and a deterministic model for tethered enzymatic SPR that are based on the same reactions. In this section we describe the reactions in general before the two models are described in the sections that follow.

The models are initialised with phosphorylated substrate distributed randomly in space (state *A*). A phosphorylated substrate can be bound by enzyme (state *B*) with first-order kinetics,

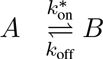

Where 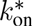 and *k*_off_ have units of s^−1^. The bimolecular on-rate (*k*_on_ with units of *μ*M^−1^s^−1^) is related to the first-order on-rate by 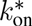 = *k*_on_[SHP-1], where [SHP-1] is the concentration of injected enzyme (in units of *μ*M). When the enzyme is bound, it can dephosphorylate substrates within reach,

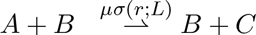

Where *C* is unphosphorylated substrate, *σ*(*r*) is the local concentration (in units of *μ*M, see above), and *μ* is the surface catalytic rate (in units of *μ*M^−1^s^−1^). We note that *μ* = 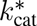 (tethered) and is used for clarity in the derivations of the mathematical models below. Lastly, phosphorylated substrate can be dephosphorylated by enzyme directly from solution,

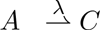

Where λ is the solution dephosphorylation rate (in units of s^−1^). The solution catalytic rate, 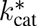 (solution) with units of *μ*M^−1^s^−1^, is related to the solution dephosphorylation rate by λ = 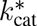 (solution)[SHP-1].

Note that *A*, *B*, and *C* represent peptide polymers that are anchored at a fixed location within the volume of the dextran matrix. We assume that the dextran matrix is stiff compared to the peptide polymers so that interactions between *A* and *B* in the matrix are determined primarily by the combined reach of the peptide polymers and the enzyme.

### Stochastic simulation

The overall state of the stochastic model can be represented by the positions of the substrate molecules, and each molecule’s current chemical state (one of *A*, *B*, or *C*). As the substrates are immobile, the system can be modeled by a collection of discrete-state jump Markov Processes with rates (i.e. propensities) for reactions as given in the preceding section. Our stochastic simulation engine generated exact realisations of these processes using the Gibson-Bruck Next Reaction Method (33) variant of the well-known stochastic simulation algorithm (SSA) (34).

The simulation is initialised with a random distribution of peptide substrates in a cube. The side length of the cube is determined by the initial concentration of peptides and the absolute number of peptides, which is a simulation parameter taken to be 500,000. For computational efficiency, we define a maximum support of 4.5 ×*L* so that reactions between a bound enzyme (*B*) and a free phosphorylated peptide substrate (*A*) that are anchored a distance larger than the maximum support are ignored. This is reasonable because the concentration of substrate that a bound enzyme experiences at the maximum support is *σ*(4.5*L*) ≈ 10^−14^ *μ*M. Increasing the maximum support produced identical simulations but required longer computational times (not shown).

### Deterministic MPDPDE calculations

As discussed in the main text, a standard model of partial-differential-equations (PDEs) that does not account for stochastic fluctuations failed to fit the tethered enzymatic SPR data and moreover, did not agree with the stochastic simulations (not shown). The low copy number of substrates within reach of tethered enzymes means that stochastic effects are prevalent. To capture these effects, we use the multi-centre particle density formalism previously used in the study of solid state physics (21, 22).

We define *ρ_m_,_m′_* as the multi-centre particle density (MPD),

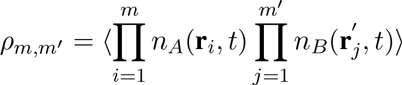

Where bold-face denotes vector quantities. The explicit expression for the first 5 MPDs are,

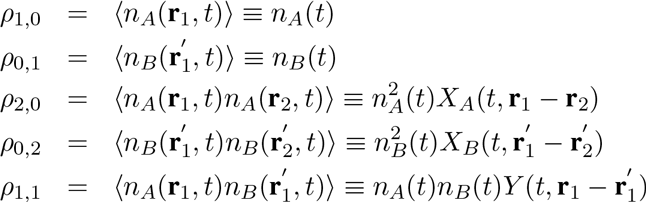

Where we have defined *n*_*A*_ and *n*_*B*_ as the concentration of *A* and *B*, respectively and *X*_*A*_, *X*_*B*_, and *Y* as the autocorrelation function for *A*, the autocorrelation function for *B*, and the pair correlation function between *A* and *B*, respectively. Note that *X*_*A*_, *X*_*B*_, and *Y* are dimensionless. The general set of PDEs governing the dynamics of the MPDs based on the reactions outlined above are,

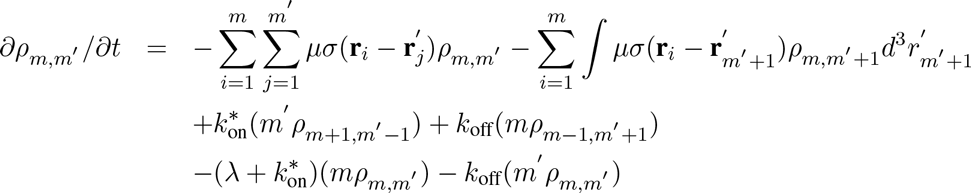

Where the parameters have been previously defined. The explicit expression for the first 5 MPDPDEs are,

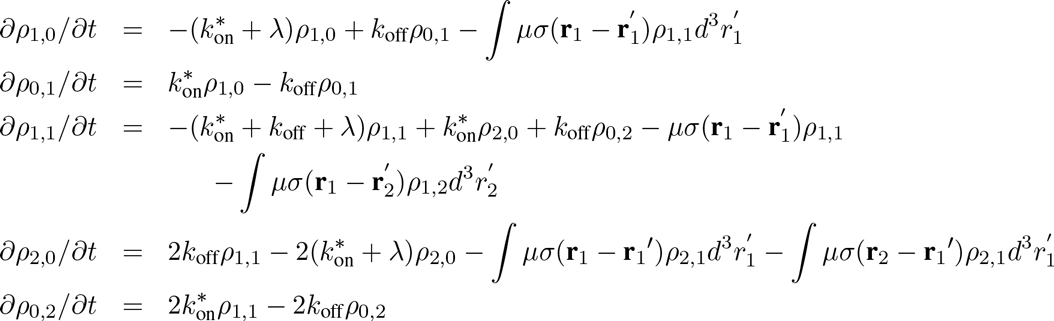

To uncouple the infinite hierarchy of these PDEs, we use Kirkwood’s approximation,

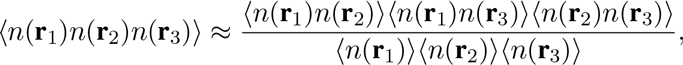

Which in our case leads to,

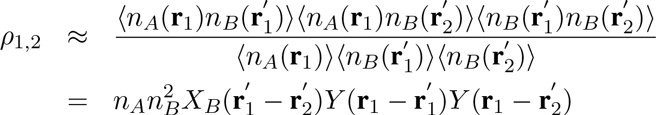

and,

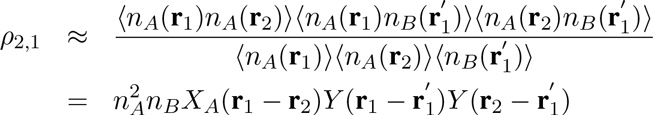

We next express the derivatives of the first five MPDs in terms of their definitions (*n*_*A*_, *n*_*B*_, *X*_*A*_, *X*_*B*_, and *Y*) to obtain,

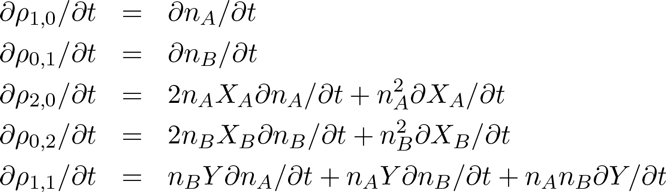

Using these derivatives together with the simplified expressions for *ρ*_1,2_ and *ρ*_2,1_ obtained using Kirkwood’s approximation, we can simplify the first 5 MPDPDEs as follows,

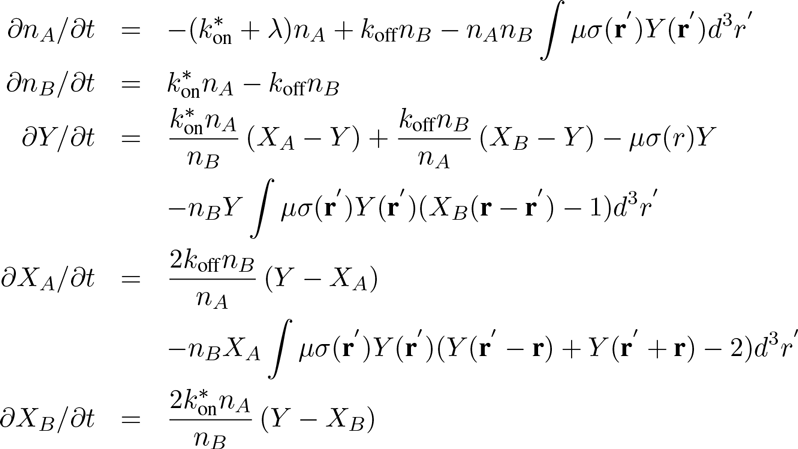

The initial conditions for this integral MPDPDE system are *n*_*A*_(*t* = 0) =[Peptide], *n*_*B*_(*t* = 0) = 0, *X*_*A*_(*t* = 0, *r*) = 1, *X*_*B*_(*t* = 0, *r*) = 1, and *Y*(*t* = 0, *r*) = 1.

A numerical solution of this integral MPDPDE system can be obtained by noting that there are two distinct types of integrals. The first integral, appearing in the equation for *n*_*A*_, is evaluated by defining |**r**′| = *r*′ to obtain,

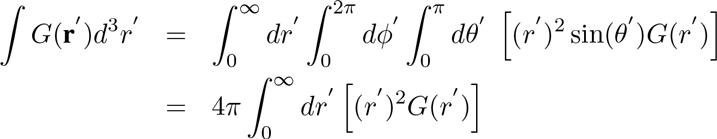

The second integral, appearing in the equations for *Y* and *X*_*A*_, is evaluated by defining *q* = |**r** ‒ **r**′ | = 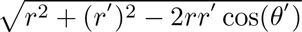 where, without loss of generality, it is assumed that **r** = r**z**̂, so that

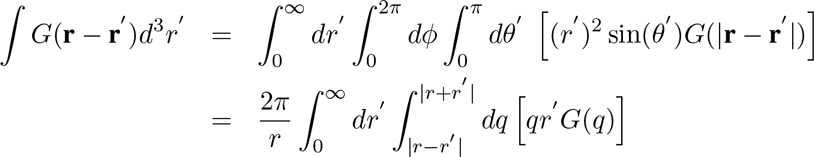

Using these integral definitions, the definition of *σ*(*r*), and by rescaling *n*_*A*_ and *n*_*B*_ by [Peptide] and r by *L*, we arrive at the following non-dimensional MPDPDE system,

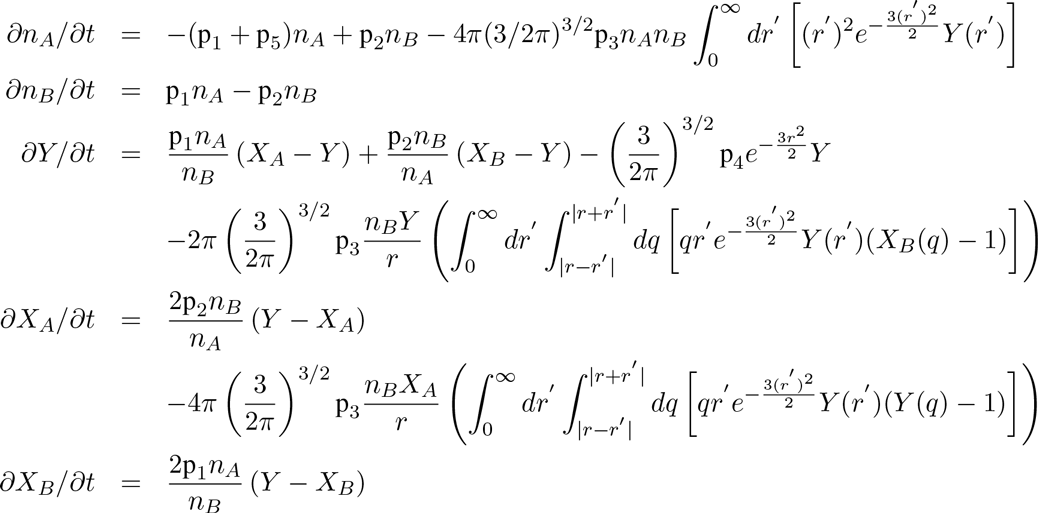

With initial conditions *n*_*A*_(*t* = 0) = 1, *n*_*B*_(*t* = 0) = 0, *X*_*A*_(*t* = 0, *r*) = 1, *X*_*B*_(*t* = 0, *r*) = 1, and *Y*(*t* = 0, *r*) = 1.

The 5 fitting parameters (p_1_, p_2_, p_3_, p_4_, and p_5_) are related to the 5 biophysical/biochemical constants as follows: p_1_ = *k*_on_ [SHP-1], p_2_ = k_off_, p_3_ = 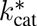 (tethered) × [Peptide], p_4_ = 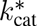 (tethered)/*L*^3^, and p_5_ = 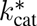 (solution) × [SHP-1].

The two numerical parameters are the spatial discretisation (ΔR) and the integration upper bound (R_max_). We found that ΔR = 0.05 and R_max_=4.5 (maximum support at which the infinite integrals were truncated) introduced errors that were substantially smaller than experimental noise while maintaining the computational efficiency required for data fitting.

### Fitting of SPR data using the MPDPDE model

The MPDPDE model was fit to the experimental data using *lsqcurvefit* in Matlab (Mathworks, MA). Specifically, the value of *n*_*B*_ from the MPDPDE model was fit to the experimental SPR data that was normalised to maximum binding. Each fit was repeated multiple times with different initial guesses for the 5 fitting parameters to make certain that the best fit was achieved (global convergence). Furthermore, we performed Markov-Chain Monte-Carlo (MCMC), using a previously published Matlab toolbox (35), on a subset of the experimental data to show that the 5 fitted parameters can be determined independently from a single SPR time course (see Supplementary Fig. 3 for MCMC analysis of the fit in Fig. 2a).

### Solution phosphatase assay

Purified SHP-1 was mixed with PEG12-ITIM peptide at 10 *μ*M, in 10 mM HEPES (pH 7.4), 150 mM NaCl, 3 mM EDTA, 0.005% Surfactant P20 and 1 mM dithiothreitol (Sigma). Temperature was regulated to 10°C with a thermocycler heat block and the production of inorganic phosphate was measured at indicated the timepoints using BIOMOL Green (Enzo Life Sciences).

The resulting progress curves (Fig. 1b) were fit using a standard mathematical model (19) based on the following reaction scheme,

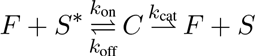

Where *F* is the phosphatase, *S*\* is the phosphorylated peptide substrate, *S* is the unphosphorylated peptide product, and *C* is the intermediate enzyme-substrate complex. This reaction scheme corresponds to the following coupled ordinary differential equations (ODEs),

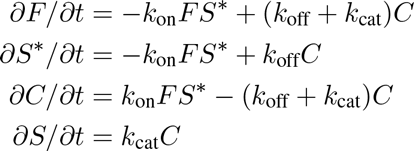

With the following conservation equations for the enzyme and substrate,

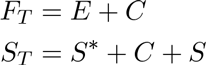

Where *F*_*T*_ is the initial SHP-1 concentration and *S*_*T*_ is the initial concentration of phosphorylated substrate. A common assumption for *in vitro* solution-based enzyme assays is that the enzyme-substrate complex (*C*) changes on a slower timescale compared to the timescale of product formation (i.e. ∂*C*/∂*t* ≈ 0). Using this quasisteady-state (QSS) approximation, which is valid when *E* ≪ *S* + *K*_*m*_ (36, 37), we find *C* = *F*_*T*_*S*\*/(*K*_*m*_ + *S*\*). Using this result, together with the conservation of substrate, we arrive at a simple ODE for the production of unphosphorylated peptide,

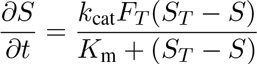

Where *k*_cat_ is the catalytic rate (in units of s^−1^) and *K*_m_ is the Michaelis-Menten constant (in units of *μ*M). The initial condition is *S*(*t* = 0) = 0.

This ODE was solved using *ode45* and fit to experimental data using the function *lsqcurvefit* in Matlab (Mathworks, MA). We found that a simultaneous fit of the model to all the data was sufficient to uniquely determine the 3 model parameters (*k*_cat_, *K*_m_, and *S_T_*).

## Acknowledgements.

We are grateful to P. Anton van der Merwe, David Vaux, Marion H. Brown, and Simon J. Davis for help with protein production, surface plasmon resonance, and/or feedback on the manuscript. We thank Natalie Haley, Jonathan Hadida, Philip K. Maini, and Vladimir Kuzovkov for feedback on the mathematical model. The work is funded by a Sir Henry Dale Fellowship (to OD) jointly funded by the Wellcome Trust and the Royal Society (Grant Number: 098363). JG was supported by an Oxford Nuffield Medical Fellowship. SAI was supported by National Science Foundation (Award Number: DMS-1255408). This work was partially supported by a grant from the Simons Foundation (to SAI). This work benefited from the Isaac Newton Institute of Mathematical Sciences (Cambridge, UK) programme on Stochastic Dynamical Systems in Biology.

## Author contributions.

JG and OD designed the research. JG, NCG, MB performed experiments. SAI and JA developed the stochastic simulation. CSS and OD developed the deterministic model and fitted data. JG, NCG, MB, JA, and OD analysed data. JG and OD wrote the manuscript. All authors read and commented on the manuscript.

## References

1. Weston CR, Davis RJ (2001) Signal transduction: signaling specificity-a complex affair. Science 292:2439–2440.

2. Shaw AS, Filbert EL (2009) Scaffold proteins and immune-cell signalling. Nature Reviews Immunology 9:47–56.

3. Garcia-Parajo MF, Cambi A, Torreno-Pina JA, Thompson N, Jacobson K (2014) Nanoclustering as a dominant feature of plasma membrane organization. Journal of Cell Science 127:4995–5005.

4. Dushek O, Goyette J, van der Merwe PA (2012) Non-catalytic tyrosine-phosphorylated receptors. Immunological Reviews 250:258–276.

5. Pei D, Lorenz U, Klingmuller U, Neel BG, Walsh CT (1994) Intramolecular regulation of protein tyrosine phosphatase SH-PTP1: a new function for Src homology 2 domains. Biochemistry 33:15483–93.

6. Pei D, Wang J, Walsh CT (1996) Differential functions of the two Src homology 2 domains in protein tyrosine phosphatase SH-PTP1. Proc Natl Acad Sci USA 93:1141–5.

7. Lorenz U (2009) SHP-1 and SHP-2 in T cells: two phosphatases functioning at many levels. Immunological reviews 228:342–59.

8. Yang J, et al. (2003) Crystal Structure of Human Protein-tyrosine Phosphatase SHP-1. Journal of Biological Chemistry 278:6516–6520.

9. Chemnitz JM, Parry RV, Nichols KE, June CH, Riley JL (2004) SHP-1 and SHP-2 associate with immunoreceptor tyrosine-based switch motif of programmed death 1 upon primary human T cell stimulation, but only receptor ligation prevents T cell activation. Journal of immunology 173:945–954.

10. Sheppard KA, et al. (2004) PD-1 inhibits T-cell receptor induced phosphorylation of the ZAP70/CD3(sig-nalosome and downstream signaling to PKC$. FEBS Letters 574:37–41.

11. Yokosuka T, et al. (2012) Programmed cell death 1 forms negative costimulatory microclusters that directly inhibit T cell receptor signaling by recruiting phosphatase SHP2. Journal of Experimental Medicine 209:1201–1217.

12. Varma R, Campi G, Yokosuka T, Saito T, Dustin ML (2006) T cell receptor-proximal signals are sustained in peripheral microclusters and terminated in the central supramolecular activation cluster. Immunity 25:117–27.

13. Yokosuka T, et al. (2008) Spatiotemporal Regulation of T Cell Costimulation by TCR-CD28 Microclusters and Protein Kinase C θ Translocation. Immunity 29:589–601.

14. Oszmiana A, et al. (2016) The Size of Activating and Inhibitory Killer Ig-like Receptor Nanoclusters Is Controlled by the Transmembrane Sequence and Affects Signaling Article. Cell Reports pp 1–16.

15. Windisch B, Bray D, Duke T (2006) Balls and chains-a mesoscopic approach to tethered protein domains. Biophysical journal 91:2383–2392.

16. Krishnamurthy VM, Semetey V, Bracher PJ, Shen N, Whitesides GM (2007) Dependence of effective molarity on linker length for an intramolecular protein-ligand system. Journal of the American Chemical Society 129:1312–1320.

17. Van Valen D, Haataja M, Phillips R (2009) Biochemistry on a leash: the roles of tether length and geometry in signal integration proteins. Biophysical journal 96:1275–92.

18. Won AP, Garbarino JE, Lim WA (2011) Recruitment interactions can override catalytic interactions in determining the functional identity of a protein kinase. Proc Natl Acad Sci USA 108:9809–14.

19. Duggleby R (1995) Analysis of enzyme progress curves by nonlinear regression. Methods in Enzymology 249.

20. Homola JJ (2008) Surface Plasmon Resonance Sensors for Detection of Chemical and Biological Species. Chemical Reviews 108:462–493.

21. Kuzovkov V, Kotomin E (1983) Some problems of recombination kinetics. Chemical Physics 76:479–487.

22. Schnorer H, Kuzovkov V, Blumen A (1990) Bimolecular annihilation reactions with immobile reactants. The Journal of chemical physics 92:2310–2316.

23. Kuzovkov V, Kotomin E (1988) kinetics of bimolecular reactions in condensed media: critical phenomoena and microscopic self-organisation. Rep. Prog. Phys. 51:1479–1523.

24. Myszka DG, He X, Dembo M, Morton Ta, Goldstein B (1998) Extending the range of rate constants available from BIACORE: interpreting mass transport-influenced binding data. Biophysical journal 75:583–94.

25. Goldstein B, Coombs D, He X, Pineda AR, Wofsy C (1999) The influence of transport on the kinetics of binding to surface receptors: application to cells and BIAcore. Journal of molecular recognition 12:293–9.

26. Sweeney MC, et al. (2005) Decoding Protein-Protein Interactions through Combinatorial Chemistry: Sequence Specificity of SHP-1, SHP-2, and SHIP SH2 Domains. Biochemistry 44:14932–14947.

27. Lee H, Venable RM, Mackerell AD, Pastor RW (2008) Molecular dynamics studies of polyethylene oxide and polyethylene glycol: hydrodynamic radius and shape anisotropy. Biophysical journal 95:1590–9.

28. Zhou H (2001) Loops in proteins can be modeled as worm-like chains. The Journal of Physical Chemistry B 105:6763–6766.

29. Hukelmann JL, et al. (2015) The cytotoxic T cell proteome and its shaping by the kinase mTOR. Nature Immunology 17.

30. Wang W, et al. (2011) Crystal structure of human protein tyrosine phosphatase SHP-1 in the open conformation. Journal of cellular biochemistry 112:2062–71.

31. Chen YNP, et al. (2016) Allosteric inhibition of SHP2 phosphatase inhibits cancers driven by receptor tyrosine kinases. Nature 2:1–17.

32. Karlsson R, Michaelsson A, Mattsson L (1991) Kinetic analysis of monoclonal antibody-antigen interactions with a new biosensor based analytical system. Journal of Immunological Methods 145:229–240.

33. Gibson MA, Bruck J (2000) Efficient exact stochastic simulation of chemical systems with many species and many channels. The Journal of Physical Chemistry A 104:1876–1889.

34. Gillespie DT (1977) Exact stochastic simulation of coupled chemical reactions. The Journal of Physical Chemistry 81:2340–2361.

35. Haario H, Laine M, Mira A, Saksman E (2006) DRAM: Efficient adaptive MCMC. Statistics and Computing 16:339–354.

36. Segel L, Slemrod M (1989) The quasi-steady-state assumption: a case study in perturbation. SIAM review 31:446–477.

37. Ciliberto A, Capuani F, Tyson JJ (2007) Modeling networks of coupled enzymatic reactions using the total quasi-steady state approximation. PLoS computational biology 3:e45.

